# SREBP1 regulates mitochondrial metabolism in oncogenic *KRAS* expressing NSCLC

**DOI:** 10.1101/2020.01.15.896373

**Authors:** Christian F. Ruiz, Emily D. Montal, John A. Haley, John D. Haley

## Abstract

Cancer cells require extensive metabolic reprogramming in order to provide the bioenergetics and macromolecular precursors needed to sustain a malignant phenotype. Mutant *KRAS* is a driver oncogene that is well known for its ability to regulate the ERK and PI3K signaling pathways. However, it is now appreciated that KRAS can promote tumor growth via upregulation of anabolic metabolism. We recently showed that oncogenic KRAS promotes a gene expression program of *de novo lipogenesis* in non-small cell lung cancer (NSCLC). To define the mechanism(s) responsible, we focused on the lipogenic transcription factor SREBP1. We observed that KRAS increases SREBP1 expression and genetic knockdown of SREBP1 significantly inhibited cell proliferation of mutant *KRAS*-expressing cells. Unexpectedly, lipogenesis was not significantly altered in cells subject to SREBP1 knockdown. Carbon tracing metabolic studies showed a significant decrease in oxidative phosphorylation and RNA-seq data revealed a significant decrease in mitochondrial encoded subunits of the electron transport chain (ETC). Taken together, these data support a novel role, distinct from lipogenesis, of SREBP1 on mitochondrial function in mutant *KRAS* NSCLC.

## INTRODUCTION

*KRAS* is the most frequently mutated oncogene in lung adenocarcinoma, present in up to 30% of cases (1–3). Lung cancer patients with tumors harboring *KRAS* mutations are associated with a poor prognosis and resistance to therapy (4). While there are covalent *KRAS*^*G12C*^ specific inhibitors and KRAS-SOS interaction directed therapeutics currently in clinical trials, there are currently no successful anti-KRAS therapies (5, 6). Accumulating studies have highlighted a potential for mutant *KRAS* to rewire cellular metabolism to promote tumor development. Substantial evidence shows that metabolic reprogramming is essential for tumor initiation and progression (7, 8). The “Warburg effect” describes a propensity for cancer cells to increase glucose uptake and convert the majority to of it to lactate even in the presence of oxygen (9). Originally, this increase in aerobic glycolysis displayed by cancer cells was attributed to damaged mitochondria. However, it is now appreciated that mitochondria remain functional in many tumors. In fact, mitochondrial metabolism is essential for providing the energy and precursors of protein, DNA, and lipids needed for the increased growth in cancer cells (10, 11). Despite growing evidence for altered metabolism in *KRAS* mutant NSCLC, how *KRAS* drives these changes is not clearly understood.

Sterol element binding regulatory proteins (SREBPs) are key transcription factors involved in regulating lipid homeostasis in all vertebrates (12). There are three SREBP isoforms in mammals: SREBP1a, SREBP1c, and SREBP2. SREBP1a and 1c are encoded by a single gene with alternative transcription start sites, whereas a separate gene encodes SREBP2. SREBP1c enhances expression of genes involved in fatty acid uptake and synthesis while SREBP1a enhances gene expression of all SREBP-responsive genes (12). SREBP2 preferentially facilitates expression of genes required for cholesterol synthesis although it can also enhance expression of genes involved in fatty acid synthesis through upregulation of the other SREBPs (13). Accumulating evidence suggests that SREBP1 is a critical link between oncogene signaling and metabolism in cancer (14–18). SREBP1’s ability to regulate fatty acid and cholesterol metabolism provides tumor cells with the energy, biomass, and reducing equivalents required for tumor growth and survival. Accordingly, several studies have reported on the capacity of SREBP1 to support tumor growth via increased fatty acid synthesis. For example, Guo et al. reported that SREBP1 signaling is required for survival of mutant *EGFR*-expressing glioblastoma (19). Several other studies have highlighted the importance of SREBP1 in cancers such as pancreatic, prostate, and colorectal cancers (17, 18, 20). However, the role of SREBP1 in NSCLC is not clear.

We recently showed that mutant *KRAS* promotes a transcriptional program of *de novo* lipogenesis in NSCLC (21). Furthermore, mutant *KRAS*-expressing cells and tumors were sensitized to the growth inhibitory effects of FASN inhibitors. This prompted us to determine exactly how KRAS was driving this lipogenic transcription program. Here we show a novel function for SREBP1 in mutant *KRAS*-expressing NSCLC distinct from lipogenesis. We demonstrate that mutant *KRAS* promotes SREBP1 expression via MEK1/2 signaling, and loss of SREBP1 decreases growth. Despite this reduction in growth, *de novo* lipogenesis was not significantly altered. Importantly, reduction of SREBP1 led to decreased expression of the mitochondrial encoded subunits of the ETC. This decrease in mitochondrial gene expression led to impaired mitochondrial metabolism as made evident by decreased oxidative phosphorylation. These results delineate a link between mutant *KRAS* and SREBP1 as well as highlight a novel role for SREBP1 on mitochondrial function distinct from lipogenesis in NSCLC.

## RESULTS

### Oncogenic KRAS increases SREBP1 expression

Mutant *KRAS* promotes a gene expression program, which drives *de novo* lipogenesis in non-small cell lung cancer (21, 22). In order to identify the transcriptional mechanism(s) responsible, we examined cDNA microarray data from studies of lungs of wildtype and *Kras*^*LSLG12D*^ mice and observed a significant increase in expression of SREBP1 in lung tumors compared to normal lung (21). To confirm the effect of mutant *KRAS* on SREBP1, we transfected HEK 293T cells with full-length (FL) SREBP1 and increasing doses of mutant *KRAS*^*G12V*^. *KRAS*^*G12V*^ induced a dose dependent increase in SREBP1 protein expression (**Figure 1a, left panel**). Classically, SREBP1 is retained in the endoplasmic reticulum. Activation of lipid sensing programs promote the cleavage of SREBP1 (cleaved SREBP1) which can then travel to the nucleus to activate transcription of its lipogenic targets. Interestingly, cleaved SREBP1 protein levels did not increase relative to full-length SREBP1 levels (**Figure 1a, right panel**) suggesting mutant *KRAS* is not affecting SREBP1 cleavage. To assess whether this was a mutant *KRAS*-dependent effect, we transfected 293T cells with SREBP1 and equal amounts of either *KRAS*^*WT*^ or *KRAS^G12V^*. While *KRAS*^*WT*^ induced SREBP1 expression slightly, the effect was much less pronounced than with *KRAS*^*G12V*^ (**Supplemental Figure 1a**).

**Figure 1:**
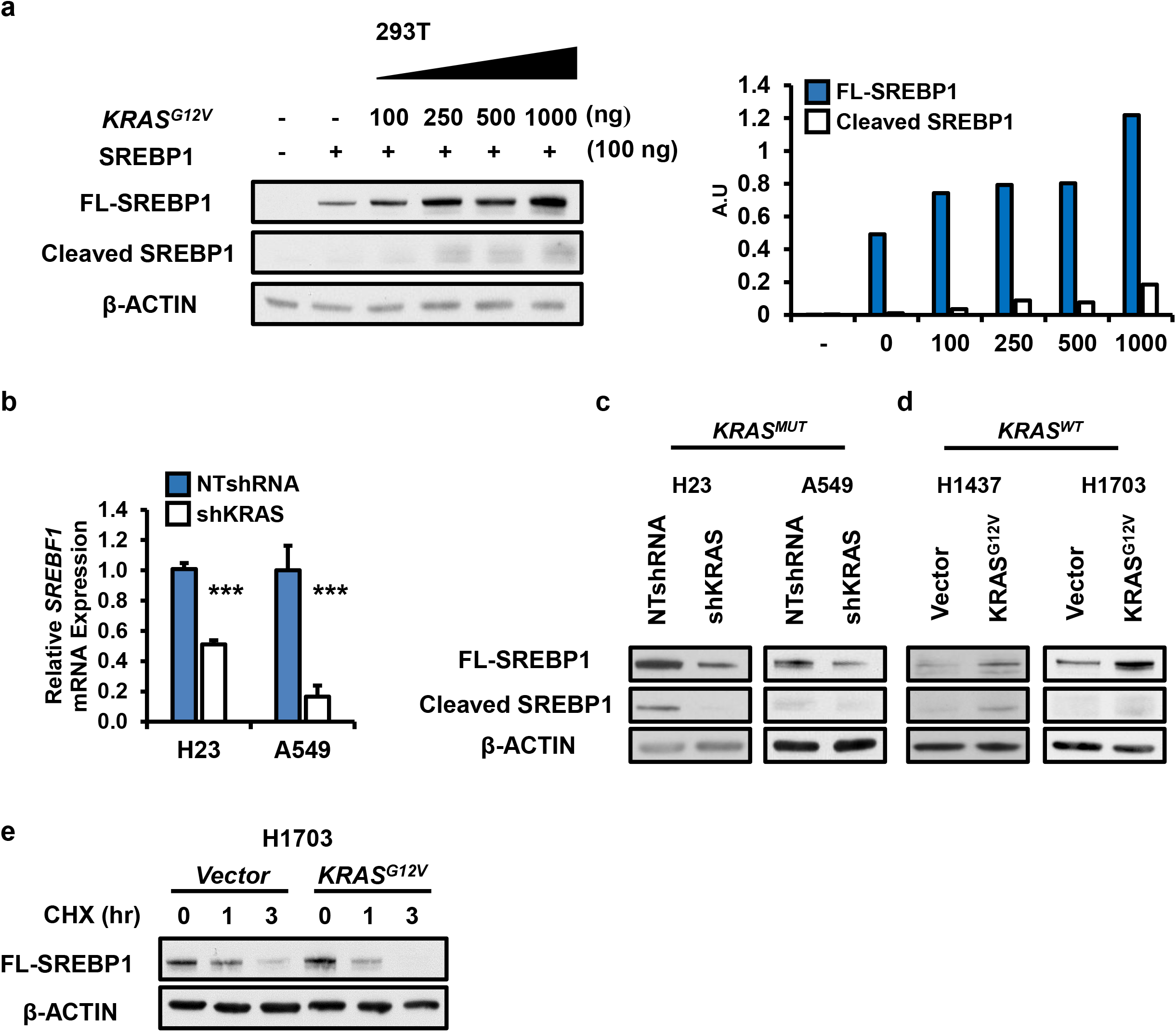
Oncogenic KRAS increases SREBP1: a) Protein expression for 293T cells transfected with full length (FL) SREBP1 in the presence or absence of increasing amounts of *KRAS^G12V^*. Protein was harvested 48 hours after transfection and analyzed via western blotting for SREBP1 and loading control, β-ACTIN. Densitometry analysis of FL-SREBP1 vs cleaved SREBP1 protein increase. Blots were analyzed using ImageJ. Values for SREBP1 were normalized to ACTIN values. SREBP1 expression for b) mRNA and c) protein in H23 and A549 stably expressing either NTshRNA or shKRAS. N=3 per group. Bars represent mean ± SD.*** p < 0.001. d) Protein expression for FL-SREBP1 and cleaved SREBP1 in H1437 and H1703 expressing either control vector, or KRAS^*G12V*^. Proteins were analyzed via western blotting. e) Protein expression for H1703 cells treated with vehicle control or 10 μM of cyclohexamide (CHX) for 1 or 3 hours (hr). Cells were expressing either vector control or *KRAS^G12V^*. Proteins were analyzed via western blotting.

Next, we investigated whether oncogenic KRAS was necessary to induce SREBP1 expression in NSCLC cells. Knockdown of KRAS in *KRAS* mutant H23 and A549 cells (1, 23) led to a reduction in SREBP1 gene expression (gene name: *SREBF1*) compared to non-target (NT) controls (**Figure 1b**). SREBP1 protein levels were similarly reduced (**Figure 1c**). In order to determine if KRAS^MUT^ was sufficient to induce SREBP1 expression, we ectopically over-expressed exogenous *KRAS*^*G12V*^ in NSCLC cells, H1437 and H1703, which are wild-type for *KRAS*. Ectopic expression of mutant *KRAS* did not significantly alter SREBP1 mRNA expression (**Supplemental Figure 1b-d**) compared to vector controls. However, endogenous SREBP1 protein expression was increased (**Figure 1d**). Similarly, in patient lung adenocarcinomas (TCGA Provisional) KRAS mutation with significantly correlated with SREBF1 mRNA abundance (p=0.03; 566 samples).

In order to identify which SREBP1 isoform (SREBP1a or SREBP1c) is predominately expressed in our NSCLC cell lines, we *in-vitro* translated SREBP1a and SREBP1c from plasmid DNA and subjected the products to SDS-PAGE in parallel with protein lysates for H23, A549, H137, H1703 (**Supplemental Figure 1e**). SREBP1a is 24 amino acids bigger than SREBP1c and they can be distinguished by size on a western blot. SREBP1c (hereafter referred to as simply SREBP1) appeared to be the predominant isoform in all four NSCLC cell lines.

Given that ectopic-overexpression of *KRAS*^*G12V*^ was not able to increase SREBP1 mRNA levels, we wanted to determine whether mutant KRAS was regulating SREBP1 expression post-transcriptionally. We inhibited protein synthesis using cycloheximide (CHX) in H1703 cells expressing either mutant KRAS (*KRAS^G12V^*) or empty vector (Vector). CHX treatment of H1703 decreased SREBP1 protein levels to a greater degree in H1703 cells expressing *KRAS*^*G12V*^ (**Figure 1e**). Together these results suggest that oncogenic KRAS increases SREBP1 protein translation in NSCLC.

### Loss of SREBP1 decreases cell proliferation in mutant*KRAS* NSCLC

Next, we sought to determine the role of SREBP1 on cell expansion in NSCLC. We generated *KRAS* mutant and *KRAS* wild-type NSCLC cell lines, which stably expressing lentiviral based NTshRNA or one of two different shRNAs targeting SREBP1, shSREBP1 # 1 and shSREBP1 # 2. We confirmed reduced expression of full length SREBP1 in shSREBP1-expressing cells by western blot (**Figure 2a, left panel**). Strikingly, loss of SREBP1 resulted in a marked reduction in cell proliferation of *KRAS* mutant cells. (**Figure 2b-c**). In contrast to mutant *KRAS*-expressing cells, SREBP1 knockdown (**Figure 2c**) had no effect or enhanced the proliferation of *KRAS^WT^*-expressing cells (**Figure 2e and 2f**). These data suggest that SREBP1 plays an essential role in the proliferation of mutant *KRAS*-expressing NSCLC cells.

**Figure 2:**
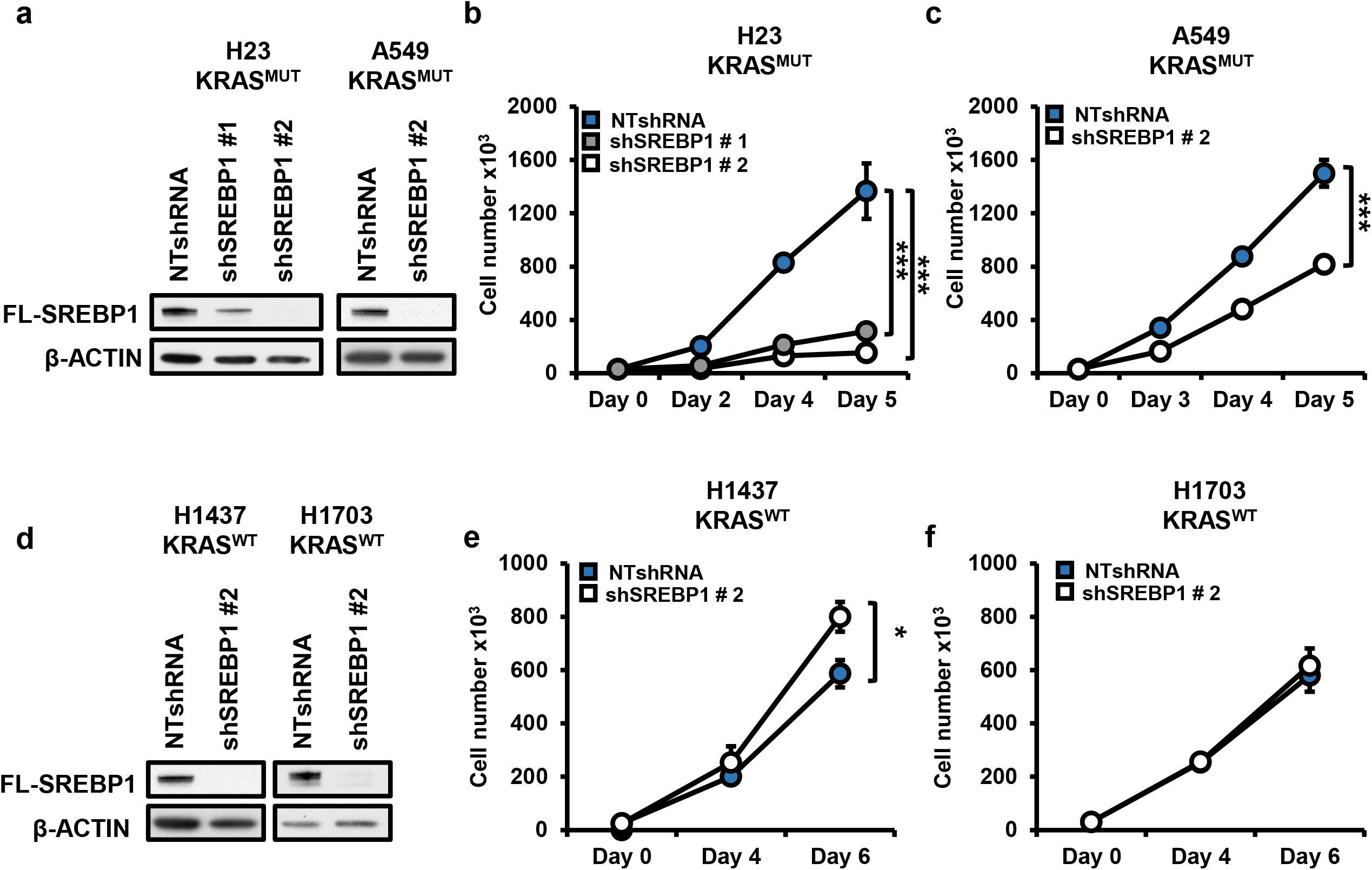
Loss of SREBP1 decreases cell proliferation in mutant *KRAS-* expressing NSCLC cells. a) SREBP1 protein expression for H23 (left), A549 (right) and f) H1437 (left), H1703 (right). Cells expressed either NTshRNA or one of two shSREBP1. Proteins were analyzed via western blot analysis. Cell number for mutant *KRAS*-expressing cells a) H23, b) A549 and *KRAS*^*WT*^ expressing cells c) H1437 d) H1703. H23 were expressing either NTshRNA or one of two shRNAs against SREBP1; shSREBP1 #1 or shSREBP1 #2. All other cells expressed either NTshRNA or shSREBP #2. Cells were seeded in 6 well plates on day 0 and counted on indicated days using an automatic cell counter. N=3 per group. Bars show ± SD. *p<0.05, ***p<0.005.

### Oncogenic KRAS regulates SREBP1 protein expression via MEK1/2 signaling

Our data demonstrate that mutant *KRAS* increases SREBP1 expression in NSCLC and in turn, SREBP1 promotes growth in mutant *KRAS*-expressing NSCLC cells. KRAS asserts many of its effects through the MEK/ERK pathway (24, 25). Previous studies showed that MEK/ERK regulate translation (26). Therefore, we treated 293T cells transfected with SREBP1 and KRAS with the MEK inhibitor AZD6244 (27–29). MEK inhibition greatly blunted the effect of KRAS on SREBP1 protein expression compared to vehicle control (**Figure 3a**). In contrast, MEK inhibition did not alter SREBP1 protein levels in 293T cells not expressing *KRAS*^*G12V*^ (**Figure 3b**). Given that MEK inhibition reduced SREBP1 protein expression in a mutant *KRAS* preferential manner, we next sought to determine whether activation of the MEK/ERK pathway alone would suffice to increase SREBP1 protein levels. We transfected 293T cells with a constitutively active *MEK1*^*D218,D222*^ allele (*MEK^DD^*) or *KRAS*^*G12V*^ (30). *MEK*^*DD*^ expression led to increased SREBP1 protein expression mimicking the effect of *KRAS*^*G12V*^ (**Figure 3c**). To further confirm that *KRAS*^*G12V*^ is regulating SREBP1 protein expression through the MEK/ERK pathway in NSCLC, we compared the effect of AZD6244 on SREBP1 levels to KRAS knockdown in H23 and A549 cells. MEK inhibitor treatment led to a significant decrease in protein levels of SREBP1 comparable to KRAS knockdown cells (**Figure 3d**). While the effect of knocking down *KRAS* on FL-SREBP1 was more striking in H23 than in A549, densitometry of A549 western blot confirms that MEK inhibition is decreasing SREBP1 expression to a greater degree in the mutant KRAS expressing cells (**Figure 3d bottom panel**). Collectively, these data strongly suggest that mutant *KRAS* increases SREBP1 protein expression via the MEK/ERK pathway.

**Figure 3:**
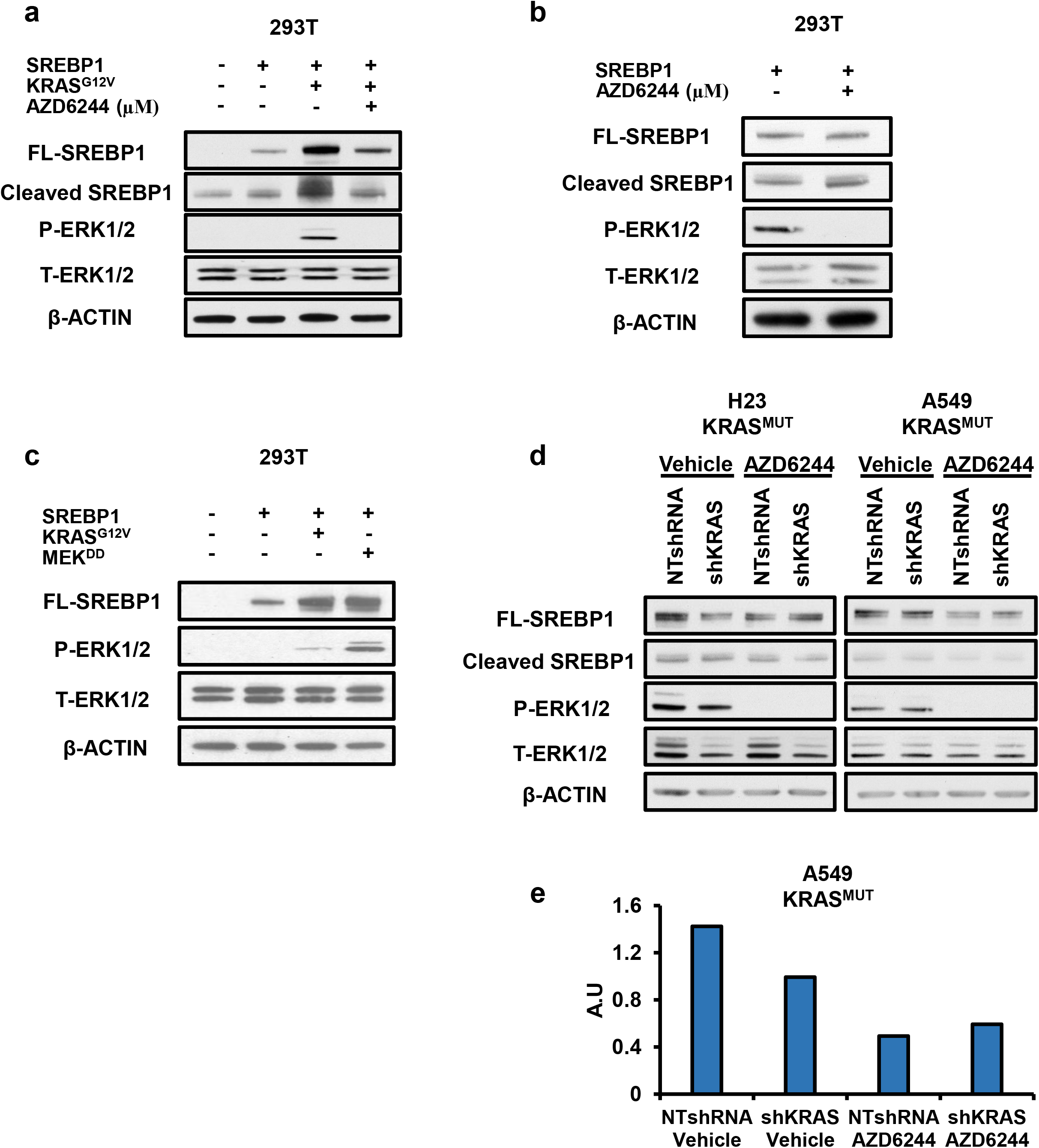
Oncogenic KRAS regulates SREBP1 protein expression via MEK1/2 signaling. a) Protein expression for 293T treated with 2.5 μM of MEK inhibitor, AZD6244, for 16 hours. 293T were transfected with either control vector or FL-SREBP1 in the presence or absence of KRAS^*G12V*^, 24 hours prior to drug treatment. b) Protein expression for 293T transfected with FL-SREBP1 and treated with AZD6244. Cells were treated 24 hours post transfections and collected 16 hours after treatment. c) Protein expression for 293T cells transfected with FL-SREBP1 and either KRAS^G12V^ or MEK^DD^. d) Protein expression for H23 (left) and A549 (right) treated with 10 μM of AZD6244 for 16 hours. Cells were stably expressing either NTshRNA or shKRAS. e) densitometry of A549 western blot from (d). FL-SREBP1 was normalized to ACTIN. Blots were analyzed using ImageJ.

### Knockdown of SREBP1 does not decrease lipogenesis in NSCLC

SREBP1 is known to induce the expression of key lipogenic enzymes including ATP Citrate lyse (ACLY), Acetyl-CoA carboxylase (ACC), and Fatty Acid Synthase (FASN), which in turn, promote cell growth by providing fatty acids, which are essential for the synthesis of membranes, energy storage, and signaling in cancer cells (12, 16, 31). Our lab has previously shown that mutant KRAS promotes the expression of these genes in NSCLC (21). We sought to determine whether SREBP1’s canonical role in lipogenesis might explain the decrease in cell proliferation observed following KRAS knockdown in KRAS^MUT^-expressing cells. We began by examining the expression of lipogenic genes in mutant *KRAS* cells with stable SREBP1 knockdown. Surprisingly, knockdown of SREBP1 did not cause a significant decrease in *ACLY*, ACACA1 (*ACC)* or *FASN* in H23 or A549 (**Figure 4a-d**). We next investigated the functional effect of SREBP1 knockdown on *de novo* lipogenesis by performing ^13^C stable isotope analysis to measure the incorporation of ^13^C glucose into palmitate, which requires ACLY, ACC and FASN (**Figure 4e**). ^13^C enrichment into palmitate was not reduced following SREBP1 knockdown, demonstrating reduced SREBP1 levels neither altered lipogenic gene expression nor lipogenesis (**Figure 4f-g**). We also measured total levels of palmitate and saw no significant difference in palmitate levels in SREBP1 knockdown cells compared to non-target controls (**Figure 4h-i**). Although SREBP2 preferentially activates expression of genes involved in cholesterol biosynthesis, it has been shown to activate expression of genes involved in fatty acid synthesis (32, 33). Importantly, we did not observe a compensatory increase in *SREBF2* levels (**Supplemental Figure 2c, left**) to rescue *de novo* lipogenesis in shSREBP1 expressing cells. Finally, *KRAS* wild-type cells subject to stable SREBP1 knockdown did not exhibit altered lipogenic gene expression or *de novo* lipogenesis (**Supplemental Figure 2**). Together, these data argue that SREBP1 maintains cell proliferation in mutant *KRAS*-expressing cells independent of its canonical role in lipogenesis.

**Figure 4:**
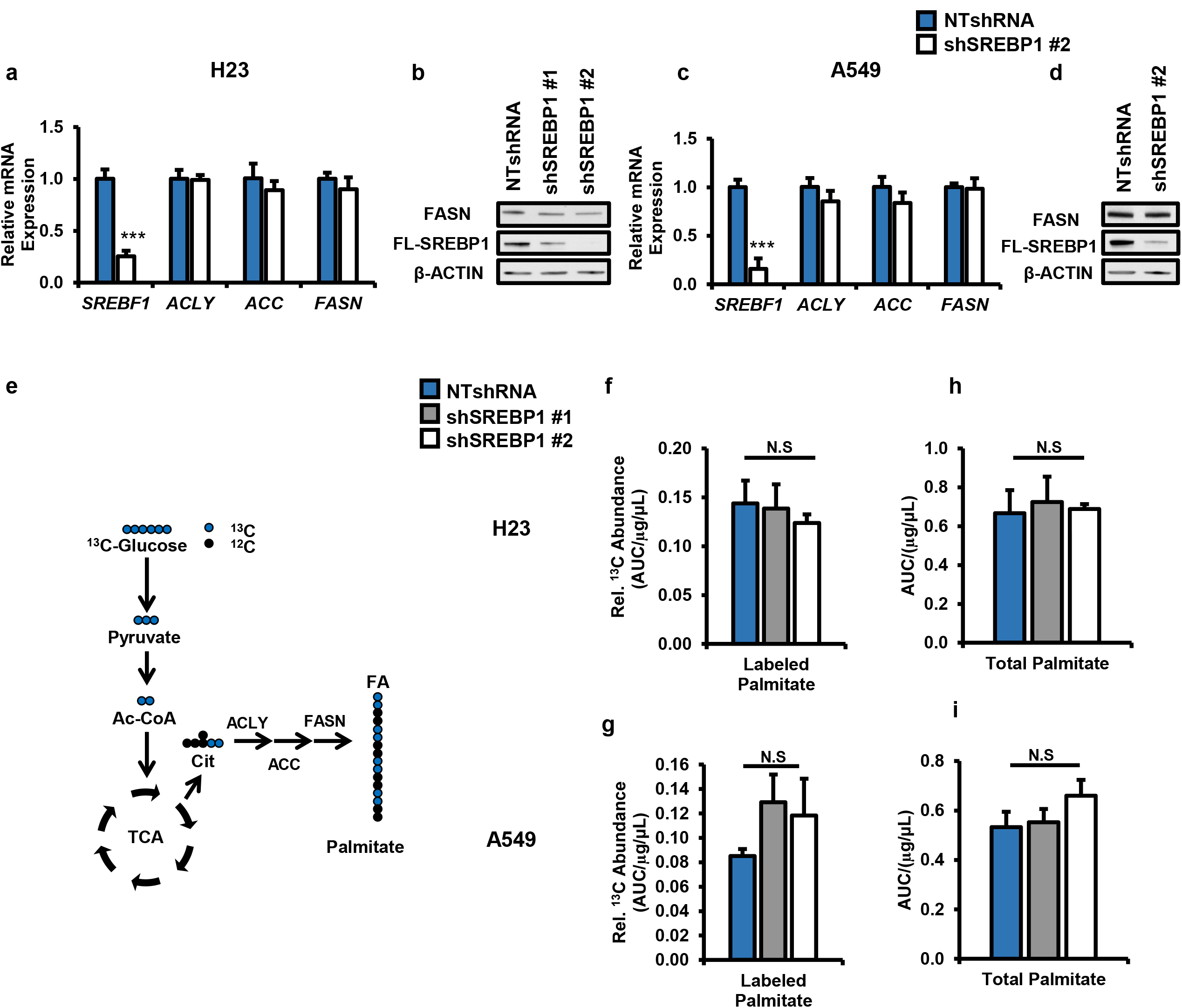
SREBP1 knockdown does not decrease*de novo* lipogenesis in mutant n *KRAS* expressing NSCLC cells. Gene and protein expression for SREBP1 and its lipogenic targets ACLY, ACC, and FASN in a) H23, b) A549. Cells expressed either NTshRNA or one of two different shSREBP1. N=3 per group. Bars indicate mean ± SD. ***P< 0.005. e) Schematic for ^13^C glucose tracer analysis on *de novo* lipogenesis. Total ^13^C glucose labeled palmitate in f) H23, g) A549. Total palmitate levels in h) H23 i) A549 cells. Palmitate was measured via GS/MS and total counts were normalized to protein concentration of cells on day of collection (μg/μl). Cells expressed NTshRNA or one of two different shSREBP1 for all metabolite tracing experiments. N=5 per group. Bars indicate mean ± SD. * p <0.05, ***P< 0.005.

### Loss of SREBP1 decreases mitochondrial-encoded electron transport chain (ETC) genes in mutant KRAS cells

The lack of changes to lipogenic gene expression in SREBP1 knockdown cells prompted us to performed RNA-seq analysis in NTshRNA and SREBP1 knockdown cells. Strikingly, we observed significant decreases in mitochondrial-encoded – but not nuclear-encoded – electron transport chain (ETC) genes (**Figure 5a**) and confirmed these findings by qRT-PCR (**Figure 5b-e**). Geneset enrichment analysis (GSEA) showed significant association (q<0.001) with TCA cycle and oxidative phosphorylation signatures (data not shown). Protein levels for mitochondrial-encoded cytochrome c oxidase I (MT-CO1) were also reduced in H23 and A549 cells expressing shSREBP1, whereas protein levels for nuclear-encoded ATP5A did not change (**Figure 5f**). In contrast, SREBP1 knockdown in KRAS^WT^ cells resulted in only minor declines to mitochondrial-encoded ETC gene expression in H1437 (**Figure 5g**) and significant increases in H1703 (**Figure 5h**). This suggests that SREBP1’s effect on mitochondrial gene expression might be facilitated by mutant *KRAS*.

**Figure 5:**
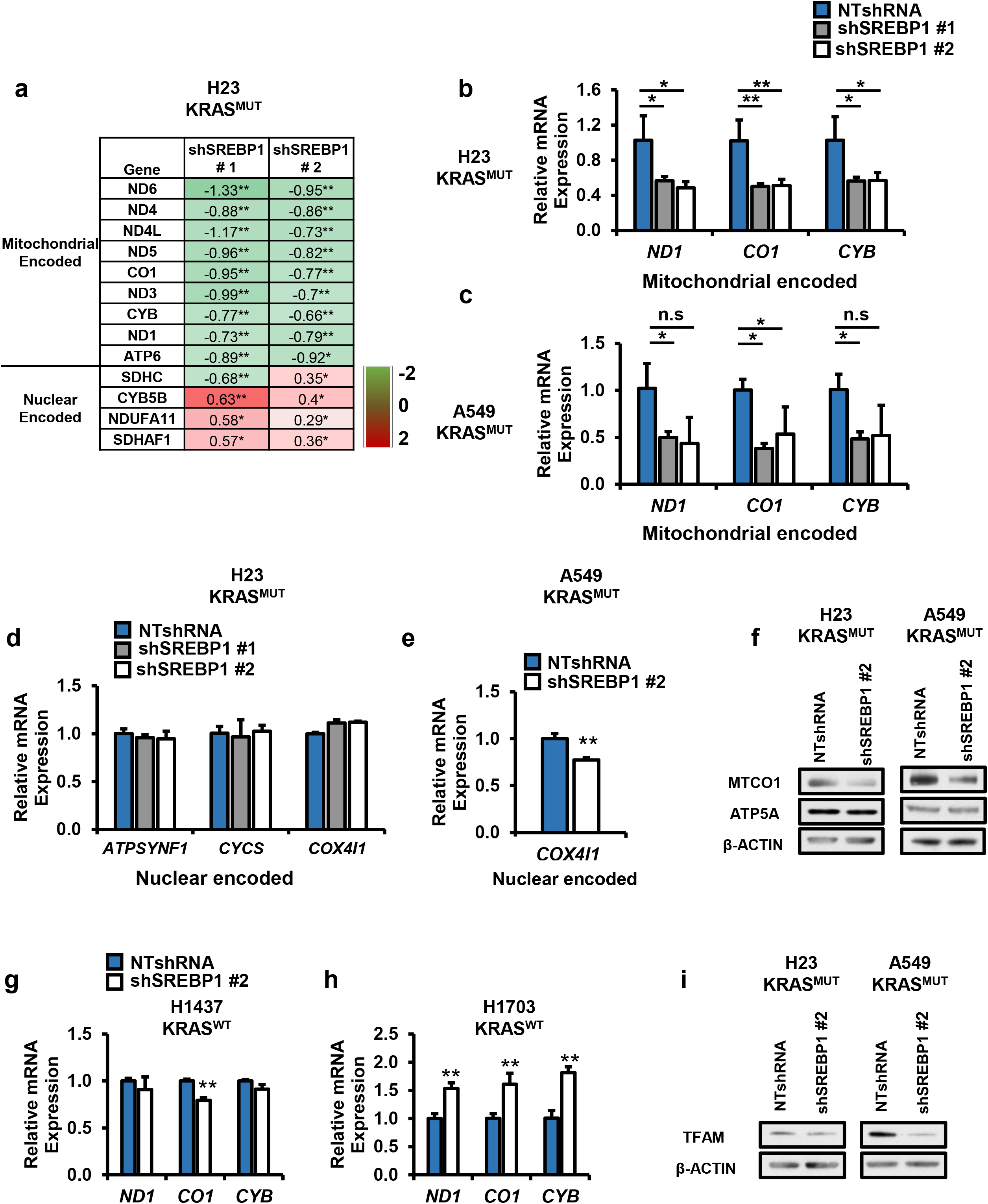
Loss of SREBP1 decreases expression of mitochondrial encoded ETC genes in mutant *KRAS* NSCLC cells. a) RNA seq analysis for mitochondrial encoded and nuclear encoded mitochondrial proteins in H23 cells expressing either NTshRNA, shSREBP1 #1 or shSREBP1 #2. shSREBP1 values were normalized to their NTshRNA expressing controls. N=3 per group. ± SD.*q <0.05 **q<0.01. Gene for mitochondrial encoded ETC genes in b) H23, and c) A549 cells expressing either NTshRNA shSREBP1 #1 or shSREBP1 #2. N=3 per group. Bars indicate ±SD. *p< 0.05. Gene expression for nuclear encoded ETC genes in d) H23 and e) A549 cells expressing either NTshRNA or one of two different shSREBP1. N=3 per group. Bars indicate ±SD. *p< 0.05. f) Protein expression for H23 (left) and A549 (right) cells expressing either NTshRNA or shSREBP1 #2. Proteins were analyzed via western blotting. Gene expression for mitochondrial encoded genes in *KRAS*^*WT*^ g) H1437 and h) H1703 cells expressing either NTshRNA or shSREBP1 #2. N=3 per group. Bars indicate ± SD. ** p <0.01.

### Loss of SREBP1 does not alter mitochondrial mass and slightly decreases copy number

Our data suggested that SREBP1 plays a role in mitochondrial biology specifically in *KRAS* mutant NSCLC cells. Indeed, mitochondrial transcription factor A (TFAM), which is required for mitochondrial DNA replication and transcription (34, 35) was significantly reduced following SREBP1 knockdown in H23 and A549 (**Figure 5i**). To determine if the loss of ETC gene and protein expression was due in part to a decrease in number of mitochondria, we stained cells with Mitotracker Green, a cell permeable dye which localizes and binds to mitochondria. There was no difference in GFP intensity, quantified by flow cytometry, between NTshRNA control and shSREBP1 cells (**Supplemental Figure 3a**). To further confirm that reduced SREBP1 expression is not affecting mitochondrial number, we measured mitochondrial copy number relative to nuclear DNA using RT-PCR, as previously described (36). There was a ~17% decrease in mitochondrial DNA copy number in SREBP1 knockdown cells compared to NTshRNA controls (**Supplemental Figure 3b**). Suggesting the decrease in mitochondrial gene transcription could be in part due to lower mitochondrial DNA content. Taken together, these data suggest that SREBP1 knockdown results in decreased expression of mitochondrial-encoded ETC genes and mitochondrial DNA copy number. Furthermore, there were no alterations to mitochondrial mass suggesting SREBP1 knockdown is affecting transcription not mitochondrial biogenesis. Lastly, nuclear encoded genes that make up subunits of the ETC do not significantly change, suggesting this is not due to SREBP1’s transcriptional activity in the nucleus.

### Loss of SREBP1 decreases oxidative phosphorylation in mutant*KRAS*-expressing NSCLC cells

Given the effect of SREBP1 knockdown on mitochondrial ETC gene expression, we wanted to determine the effects on mitochondrial function. Knockdown of SREBP1 resulted in a significant decrease in basal oxygen consumption rate (OCR) (~80%) and maximal respiration (~70%) (**Figure 6a-c**) compared to NTshRNA cells expressing mutant *KRAS*. In contrast, we did not see any difference in basal OCR in *KRAS*^*WT*^ cells (H1437) with stable knockdown of SREBP1 (**Supplemental Figure 4a-b**). A major fuel for oxidative phosphorylation via the TCA cycle is glucose. Therefore, we performed ^13^C glucose tracer analysis to determine whether SREBP1 knockdown alters glucose utilization by the TCA cycle. We measured enrichment of m+2 metabolites into the TCA, since they would be derived from labeled glucose (**Figure 6d**). We observed a significant decrease in m+2 citrate (~56%), fumarate (~43%), and malate (~40%) when SREBP1 was knocked down in H23 cells (**Figure 6e**). By contrast, we did not see a significant change in these metabolites in *KRAS*^*WT*^ cells (H1437) with SREBP1 knockdown (**Figure 6f**). These data suggest that knockdown of SREBP1 impairs oxidative phosphorylation from glucose in mutant *KRAS*-expressing cells.

**Figure 6:**
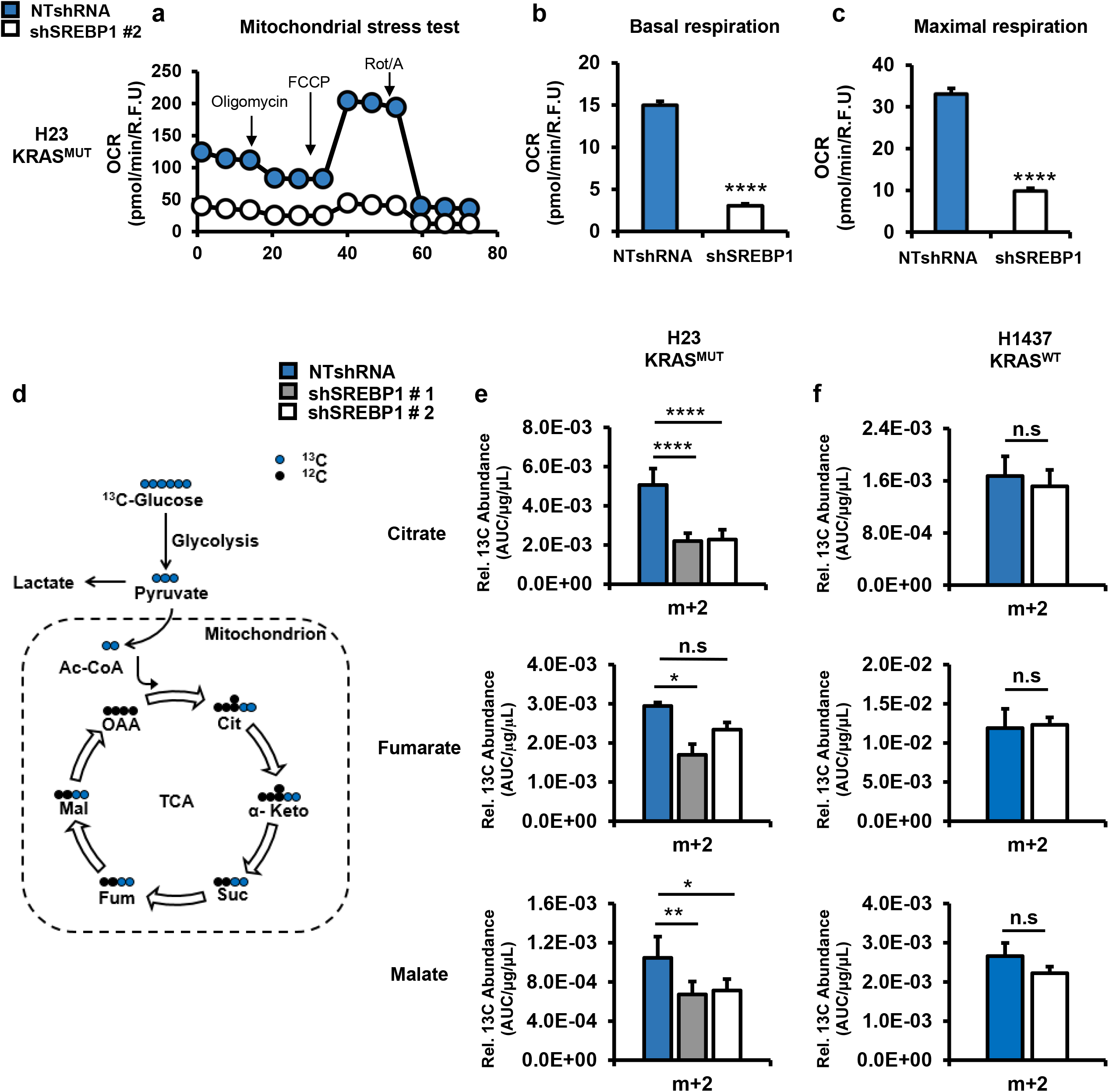
Loss of SREBP1 decreases oxidative phosphorylation in mutant *KRAS*-expressing NSCLC cells. a) Representative mitochondria stress test performed on H23 expressing NTshRNA or shSREBP1 #2. The stress test provides b) basal respiration and c) maximal respiration. OCR was measured using a Seahorse Bioenergetic Flux Analyzer. N≥10 per group. Bars indicate mean ± SE. ****p < 0.0001. d) Schematic of glucose utilization by the TCA cycle into m+2 intermediates. Relative amount of ^13^C labeled m+2 citrate, fumarate, and malate in e) H23 and f) H1437 cells. Cells expressed NTshRNA or one of two different shSREBP1. Cells were labeled with ^13^C [U6] glucose, and harvested after 6 hr, and analyzed by GC/MS for TCA cycle metabolites. N=5 per group. Bars indicate ± SD. * p <0.01, ** p <0.001.

Given the increased expression of SREBP1 in NSCLC, we examined the effect of SREBP1 transcript levels on overall survival in patients with lung adenocarcinoma using Kmplotter (37). Kmplotter is an online survival analysis software that allows for the meta-analysis of patient data from an integrative lung cancer microarray database. Overall survival was significantly lower in patients with tumors that have high expression of SREBP1 (p<0.01) (**Supplemental Figure 5**). It is important to note however that the survival curve does not consider mutant *KRAS* expression. Nonetheless, these data suggest SREBP1 expression is negatively associated with survival in patients with lung adenocarcinoma.

## DISCUSSION

Activating mutations of *KRAS* drive metabolic alterations that promote tumor growth in NSCLC (21, 22, 24, 38–41). However, the detailed molecular mechanisms by which KRAS regulates metabolism in NSCLC are not well understood. Here, we report a novel role for SREBP1 distinct from lipogenesis in *KRAS*-expressing NSCLC. Oncogenic KRAS increases SREBP1 expression and loss of SREBP1 leads to decreased cell proliferation independent of its role in lipogenesis. Importantly, high SREBP1 expression correlates with poor survival in patients with lung adenocarcinoma. Most interestingly, we report for the first time, to our knowledge, that loss of SREBP1 in mutant *KRAS*-expressing NSCLC leads to reduction of mitochondrial-encoded ETC subunits, resulting in deficient mitochondrial metabolism.

Mutant *KRAS* activates over a dozen downstream targets to assert its pro-tumorigenic effects and the RAF/MEK/ERK pathway is among one of the most well-characterized (15). In fact, many approaches to targeting mutant *KRAS* cancers involve the utilization of MEK/ERK inhibitors (27–29, 42–44). Our work revealed that KRAS regulates SREBP1 expression via MEK/ERK activation. MEK inhibition using AZD6244 greatly reduced the effect of mutant KRAS on SREBP1 protein expression in 293T cells. Furthermore, activation of MEK pathway with constitutively active *MEK1* mutant, *MEK^DD^*, was sufficient to increase SREBP1 protein expression. Similarly, MEK inhibition in NSCLC cells (H23 and A549) reduced SREBP1 levels similar to *KRAS* knockdown. Multiple ERK1/2 phosphorylation sites have been mapped on SREBP1 (45), and ERKs 1/2 are the only known targets of MEKs 1/2, implicating ERK in mutant *KRAS* mediated regulation of SREBP1. Interestingly, mutant KRAS does not appear to regulate SREBP1 cleavage, suggesting KRAS controls SREBP1 activity independent of cleavage. However, additional inhibitor studies need to be performed to fully elucidate the mechanism(s) responsible for KRAS regulation of SREBP1. Beyond characterizing ERK1/2 phosphorylation sites on SREBP1, further work should also include proteomic analysis to map out all major post-translational modifications on SREBP1 in the presence and absence of mutant *KRAS*.

Our results support the notion that SREBP1 is important for mutant *KRAS*-expressing NSCLC cell viability (16–18, 46); however we were surprised to find that loss of SREBP1 did not significantly alter gene expression of classic lipogenic targets *ACLY, ACACA1, and FASN* (21). Furthermore, using ^13^C tracer analysis with GC/MS, we found that mutant *KRAS*-expressing NSCLC with reduced levels of SREBP1 could still make sufficient levels of saturated fatty acids such as palmitate. Williams et al. showed that an essential requirement for SREBP1 is to maintain the ratio of monosaturated vs monounsaturated fatty acids (47). In their study, loss of SREBP1 did not lead to decreases in palmitate but instead a significant decrease in monounsaturated fatty acids such as oleate which ultimately resulted in lipotoxicity and cell death. However, these studies were carried out in glioma cells, which are not mutant KRAS dependent. Additionally, oleate levels did not decrease in our models when SREBP1 was knocked down (data not shown), suggesting an alternative mechanism for loss of cell proliferation.

Earlier studies focused on SREBP1’s role in lipid homeostasis and regulation via cleavage in low-cholesterol environments (12, 14, 19, 48). Recently, however, multiple studies have unraveled novel roles for SREBP1 in unexpected pathways linked to diabetes, cancer, the immune system, and autophagy (15, 49–53). Using RNA-seq analysis, we discovered loss of SREBP1 resulted in decreased mitochondrial gene expression in NSCLC cells. Loss of SREBP1 also reduced protein levels of mitochondrial transcription factor A (TFAM), which is one of three key transcription factors required for mitochondrial DNA replication and transcription (34). Interestingly, the effect of SREBP1 on TFAM appeared to be translational since we did not see a difference in TFAM transcript in our RNA-seq analysis of SREBP1-knockdown cells. This suggests that SREBP1 is playing a role downstream of TFAM transcription. Decreased mitochondrial gene expression resulted in impaired mitochondrial function characterized by reduced TCA cycle flux and oxygen consumption. It is not clear whether decreased proliferation in SREBP1 knockdown cells is due to SREBP1’s effects on the mitochondria. Furthermore, our work did not establish whether SREBP1’s effect on the mitochondria is strictly mutant *KRAS*-dependent. While mutant *KRAS* expression was sufficient to enhance SREBP1’s effect on the mitochondria as shown by genetic knockdown and overexpression experiments, further studies are required to determine to what extent KRAS is important in SREBP1-mediated mitochondrial metabolism and transcription. Additionally, it remains to be seen whether other prominent oncogenes in lung cancer, such as mutant EGFR which also activates ERK1/2, similarly alter SREBP1 function. Our results also suggest an alternative pathway for KRAS mediated lipogenesis in NSCLC since loss of SREBP1 showed no significant decrease in *de novo* lipogenesis. Further studies are required to illuminate how KRAS is regulating fatty acid synthesis which could potentially be via other lipogenic transcription factors implicated in cancer such as carbohydrate responsive element–binding protein (ChREBP) (54). Finally, the finding that SREBP1 plays a role in mitochondrial homeostasis presents a novel opportunity for targeted therapy in *KRAS* mutant lung cancers.

## Materials and Methods

### [^13^C] isotopomer analysis

Cells were seeded in 6-cm culture dishes (800,000 cells per dish) overnight. The following day, cells were washed twice with warm 1X PBS, and medium was changed to RPMI with 10 mM [U-^13^C6] glucose (Cambridge Isotopes) as the only glucose source and 10% dialyzed FBS and 2 mM glutamine for 6-16 hours.

#### Lipid extraction and GCMS analysis

Cells were harvested in 0.9% NaCl and centrifuged at 10,000 RPM at 4°C. The pellet was re-suspended in 2:1 chloroform:methanol. Before drying down under nitrogen, 50 nmoles of heptadecanoic acid was added to all samples as an internal control. Fatty acids were then saponified as previously described (21). Following saponification, metabolites were dried down under nitrogen again and methylated with boron trifluoride (Sigma, 15716). Mass spectral data were obtained on an Agilent 7890B Gas Chromatograph coupled with an Agilent 5977A MDS. The settings were as follows: GC inlet 230 °C, transfer line 280 °C, MS source 230 °C MS Quad 150 °C. An HP-5MS column (30 M length, 250 μm diameter, 0.25 μm film thickness) was used for fatty acid analysis and palmitate and its isotopomers were monitored at 270-286 *m/z*.

#### TCA cycle metabolite extraction

Intermediate metabolites were harvested in 80% methanol in water with 10 nmoles adonitol per sample as internal control. Metabolites were dried down under nitrogen and derivatized as previously described (55). In brief, cells were frozen and thawed three times, and centrifuged, and the supernatant was collected. The supernatant was then dried down and methoximated using MOX (Thermo Scientific, TS-45950) and derivatized with BSTFA (TCI, B3402). All metabolite data was analyzed using Mass Hunter and abundance corrected using ISOCOR.

### Analysis of SREBP1 expression

Publicly available data from the The Cancer Genome Atlas (TCGA) were analyzed for SREBP1 expression in 720 lung tumors. Overall survival and the hazard ratio were graphed and calculated using an online tool called KmPlotter (37). All cases analyzed were adenocarcinomas.

### Cell culture and reagents

All cells were purchased from American Type Culture Collection (Manassas, Virginia, United States) and cultured under recommended conditions. Specifically, H23, A549, H1437 and H1703 cells were cultured in Roswell Park Memorial Institute (RPMI) 1640 medium (Corning) and Beas-2B and 293T cells were cultured in Dulbecco’s Modified Eagle’s Medium (DMEM) (Corning). All media were supplemented with 10% fetal bovine serum (FBS) (Gemini) unless indicated. Delipidated media (DL) was RPMI media supplemented 10% delipidated FBS (Gemini). Cells were transfected using Attractene reagent as per manufacturer’s suggestions (QIAGEN). Plasmids used in all transfections are listed in **Table 6.2**. The cell lines were routinely tested for mycoplasma contamination (56). All cells were incubated at 37° C with 5% CO_2_.

### Genetic manipulation of KRAS*in vitro*

H1437, H1703, H858, and H1299 cells were tranduced with retrovirus expressing Kras^G12V^ (pBabe KRAS^G12V^) or either green fluorescent protein (GFP) vector control (pCMV-GFP) or empty vector (pMSCV-Puro) to serve as controls. Retrovirus was generated using CaP transfection into phoenix cell. Cells were transfected with expression vectors for vsv and gag-pol with the retroviral vector of interest. A549 and H23 NSCLC cells were transduced with lentivirus expressing non-target short hairpin RNA (NTshRNA) pLKO (NTshRNA) or shRNA against Kras (shKRAS;TRCN0000033262, MilliporeSigma, Burlington, MA, USA). Lentivrus was prepared as described above. Following infection, cells were selected in puromycin (1 μg/ml) for one week to establish stable pools.

### Genetic manipulation of SREBP1*in vitro*

For stable SREBP1 knockdown, H23, A549, H1437 and H1703 cells were infected with non- target shRNA lentivirus (NTshRNA) or lentivirus with one of two different commercially available shRNAs against SREBP1 (shSREBP1 #1:TRCN0000020605, shSREBP1 #2: TRCN0000020607) (MilliporeSigma, Burlington, MA, USA). Following infection with virus, H23, A549, and H1703 cells were selected in 1 μg/ml of puromycin. H1437 were selected in 2 μg/ml of puromycin. All cells were grown in indicated doses of puromycin for one week to establish stable pools.

### *In vitro* translation

In vitro translation of full-length SREBP1a and full-length SREBP1c was carried out using the TNT rabbit reticulocyte lysate translation kit (Promega, Wisconsin, USA) as per the manufacturer’s instructions. In brief, plasmids with the open reading frames of SREBP1c and SREBP1a were mixed separately with the components of the TNT rabbit reticulate lysate kit and incubated at 30°C for 90 mins. Following incubation, the product was diluted 1:5 in water and subjected to SDS-PAGE alongside lysates of H23, A549, H1437, and H1703 cells.

### Mitochondrial DNA copy number

Mitochondrial DNA copy number was measured as previously described (36). In brief, H23 cells expressing NTshRNA or shSREBP1 # 1 were seeded in triplicate in a 6-well culture dish (Corning). Genomic DNA was isolated using a commercially available kit (Invitrogen, K182001) as per the manufacturer’s instructions. Isolated DNA was subjected to PCR using primers for nuclear DNA (B2M) or mitochondrial DNA (ND1). The relative mitochondrial DNA content was then determined as follows:

a. ΔC_T_ = (nuclear DNA C_T_ – mito DNA C_T_)
b. Relative mitochondrial DNA content = 2 × 2^ΔCT^

### Mitochondrial oxygen consumption

Oxygen consumption rates were measured using a Seahorse Bioenergetic Flux analyzer (XFe96). Basal respiration and ATP-coupled respiration, represented as OCR, were measured using a Mitochondrial Stress Test assay as per manufacturer’s instructions (Agilent, 103015-100).

### Proliferation studies

Growth Curves: Cells were seeded into 6-well dishes (Corning) with an initial seeding density of 30,000-50,000 cells per well and counted on days indicated using a Countess Automated Cell Counter (Thermo Fisher Scientific, Waltham, MA, USA) cell automated countess (Invitrogen). For treatment with inhibitors, ROS, and nutrients, cells were treated the morning after plating and final counts were performed 3 days later.

### Real-time RT-PCR

Total RNA was extracted from tumors and cells with the RNeasy Kit (Qiagen, Hilden, Germany). The reverse-transcription reaction was performed with a high-capacity cDNA Synthesis Kit (Applied Biosystems). Real-time quantitative PCR analyses of human genes were performed, as previously described (21). All primers used are listed in **Table 6.3.** One of three housekeeping genes, 18s, HPRT, or β-ACTIN was used for normalization.

### RNAseq analysis

RNAseq was performed by Novagene (Novagene Sacramento, CA). Illumina HiSeq RNA sequencing of triplicate FASTQ file reads passing Illumina purity filter were aligned using TopHat2 and Cufflinks, with statistical analysis performed by CuffDiff, generating files of normalized counts for detected genes and transcripts (UCSC hg38). The Galaxy server at UCLA (galaxy.org) was used for FASTQ alignment and analysis (57). Aligned RNAs passing QC thresholds were used to calculate transcript abundance ratios followed by log2 linear scaling. Functional association with RNA abundance changes was assessed by gene set enrichment (GSEA). As expected, multiple signatures associated with mitochondrial function were significantly enriched with normalized enrichment score (NES) p value <0.05 and the false discovery rate (FDR) q value <0.05.

## Acknowledgements

We would like to thank Geoffrey D. Girnun, PhD, for his scientific discussions and editing of the text. We would also like to thank Alex Bott, PhD, and Mandar Muzumdar, MD, for critical editing of the manuscript. This work was supported in part by funds from the Ruth L. Kirschstein National Research Service Award (NRSA) F31 Grant 1F31CA210626-01.

---

**Supplemental Figure 1:**
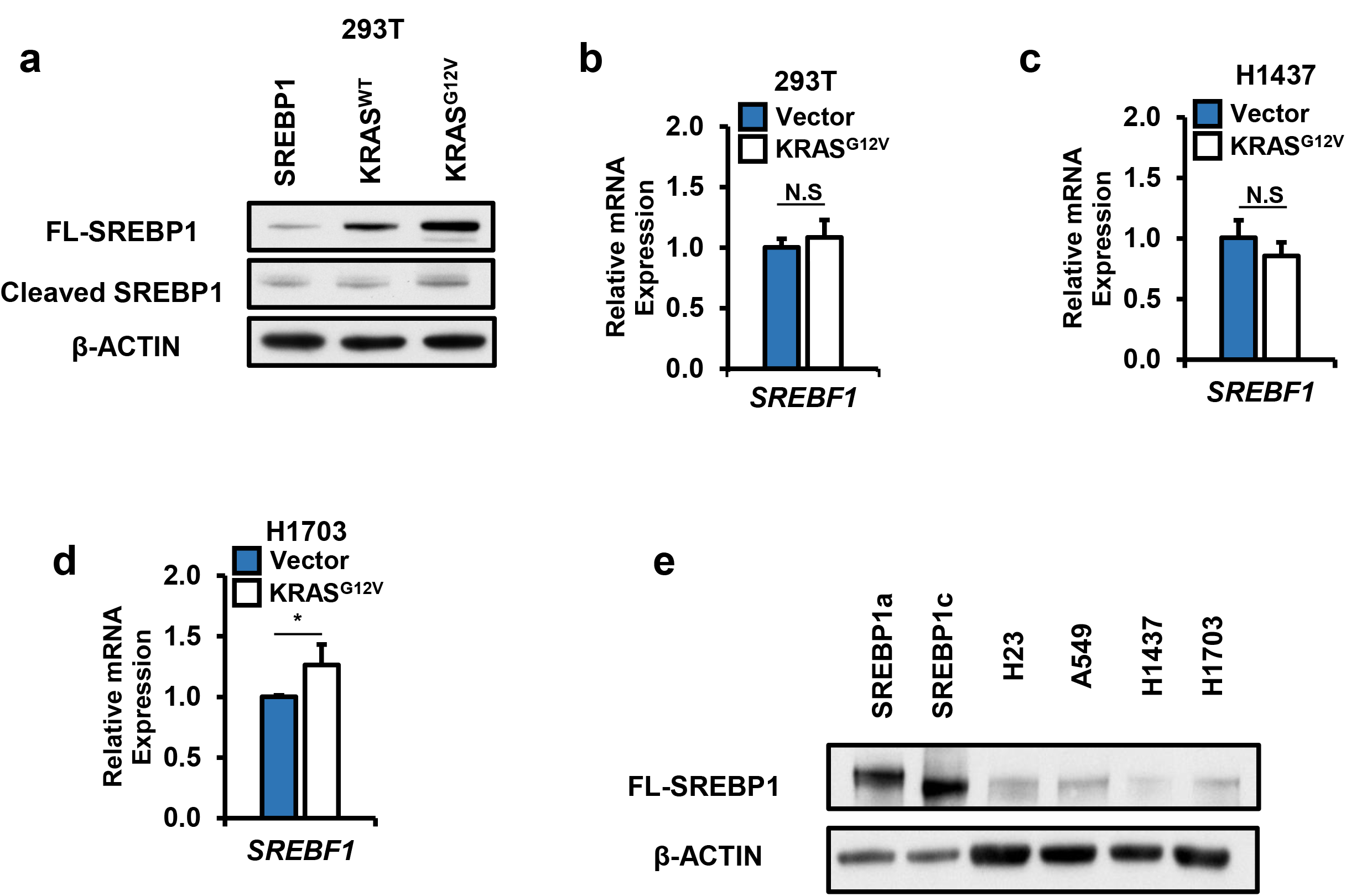
Oncogenic KRAS increases SREBP1 expression: a) Protein expression in 293T cells transfected with FL-SREBP1 in the presence of either KRAS^WT^ or KRAS^G12V^. SREBP1 mRNA expression for b) 293T, c) H1437, and d) H1703 cells. Cells were expressing empty vector (Vector) or KRAS^G12V^. N=3 per group. Bars indicate ± SD. * p <0.01 e) Full length (FL) SREBP1a and SREBP1c were produced via *in vitro* translation and subjected to SDS-PAGE alongside lysates of H23, A549, H1437, and H1703 cells.

**Supplemental Figure 2:**
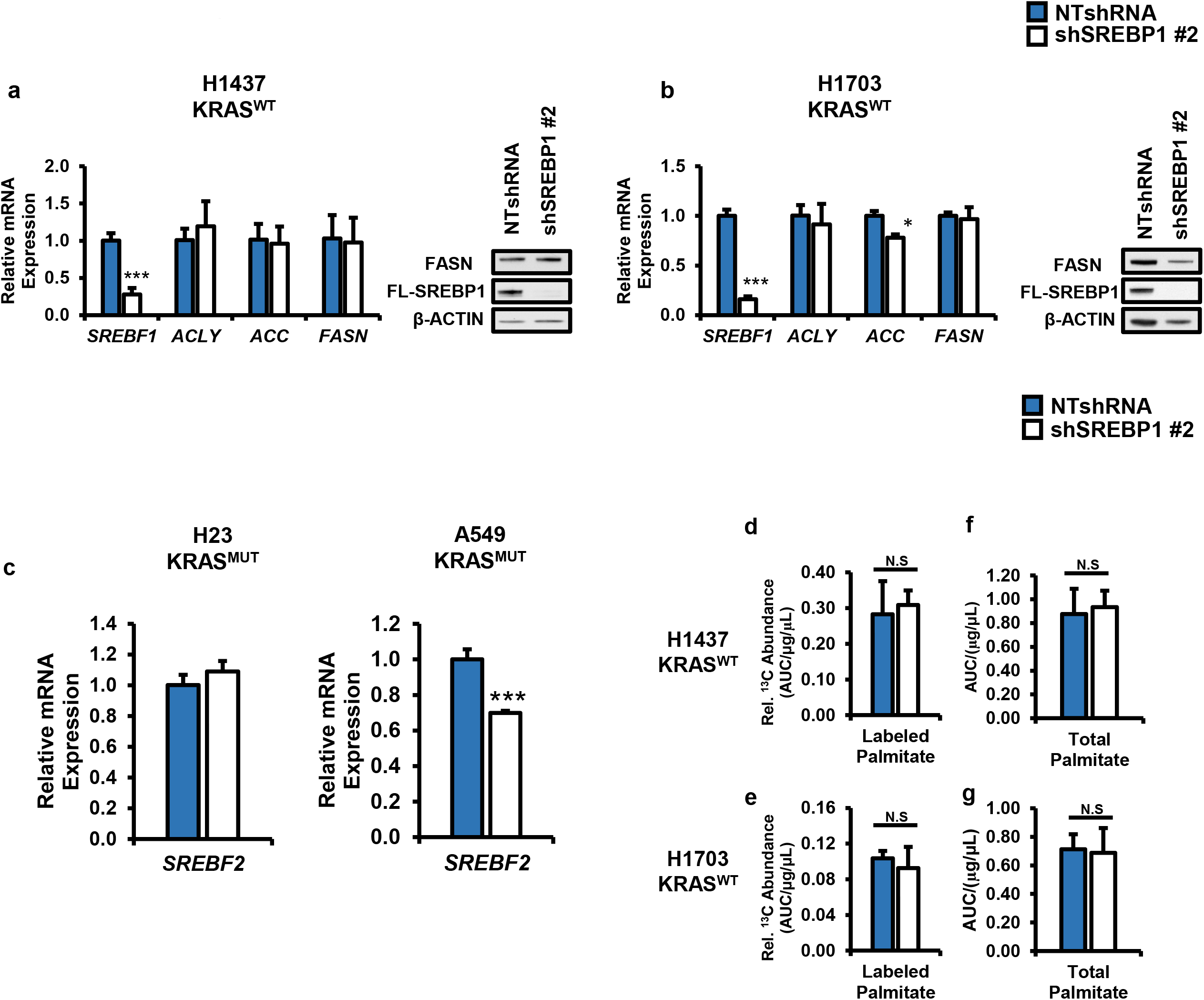
SREBP1 knockdown does not decrease lipogenesis in NSCLC. Gene and protein expression for SREBP1 and its lipogenic targets *ACLY, ACC (gene name: ACACA1), and FASN* in *KRAS*^*WT*^ expressing cells. a) H1437, b) H1703. Cells expressed either NTshRNA or shSREBP1 #2. N=3 per group. Bars indicate ± SD. c) *SREBF2* gene expression for H23 (left) and A549 (right). Total ^13^C glucose labeled palmitate in d) H1437, e) H1703. Total palmitate levels in f) H1737, and g) H1703. Palmitate was measured via GS/MS and total counts were normalized to protein concentration of cells on day of collection (μg/μl). Cells expressed NTshRNA or shSREBP1 #2 for all metabolite tracing experiments. N=5 per group. Bars indicate mean ± SD. * p <0.05, ***P< 0.005.

**Supplemental Figure 3:**
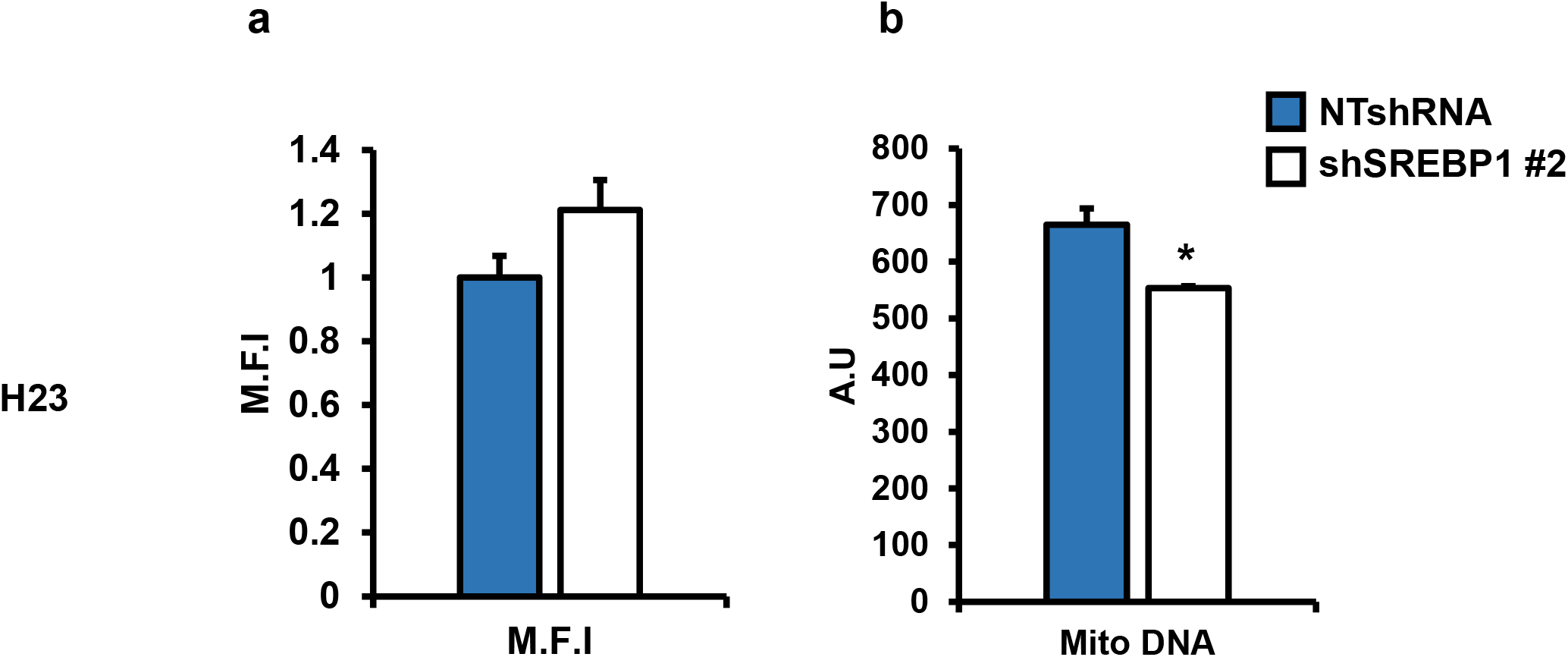
Loss of SREBP1 does not alter mitochondrial mass and slightly decreases copy number. a) Mean fluorescence intensity (M.F.I) of H23 cells stained with MitoTracker Green. Cells were stably expressing either NTshRNA or shSREBP1 #2. Fluorescence was measured via flow cytometry. b) Mitochondrial DNA content relative to nuclear DNA content in H23 expressing either NTshRNA or shSREBP1 #2. DNA was harvested and subjected to qRTPCR. AU= Abitrary units. N=3 per group. Bars indicate mean ± SD.

**Supplemental Figure 4:**
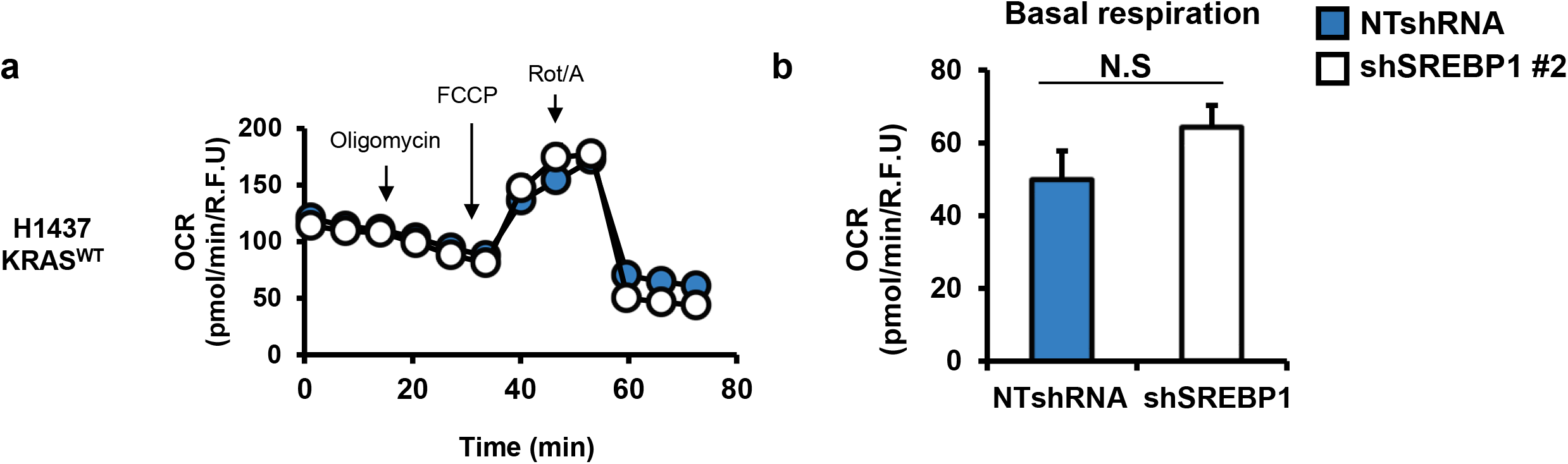
Loss of SREBP1 decreases oxidative phosphorylation in mutant*KRAS* NSCLC cells. a) Representative mitochondria stress performed on H1437 cells expressing NTshRNA or shSREBP1 #2. Test provides b) Basal OCR. N≥10 per group. Bars indicate mean ± SE. OCR was measured using a Seahorse Bioenergetic Flux Analyzer.

**Supplemental Figure 5:**
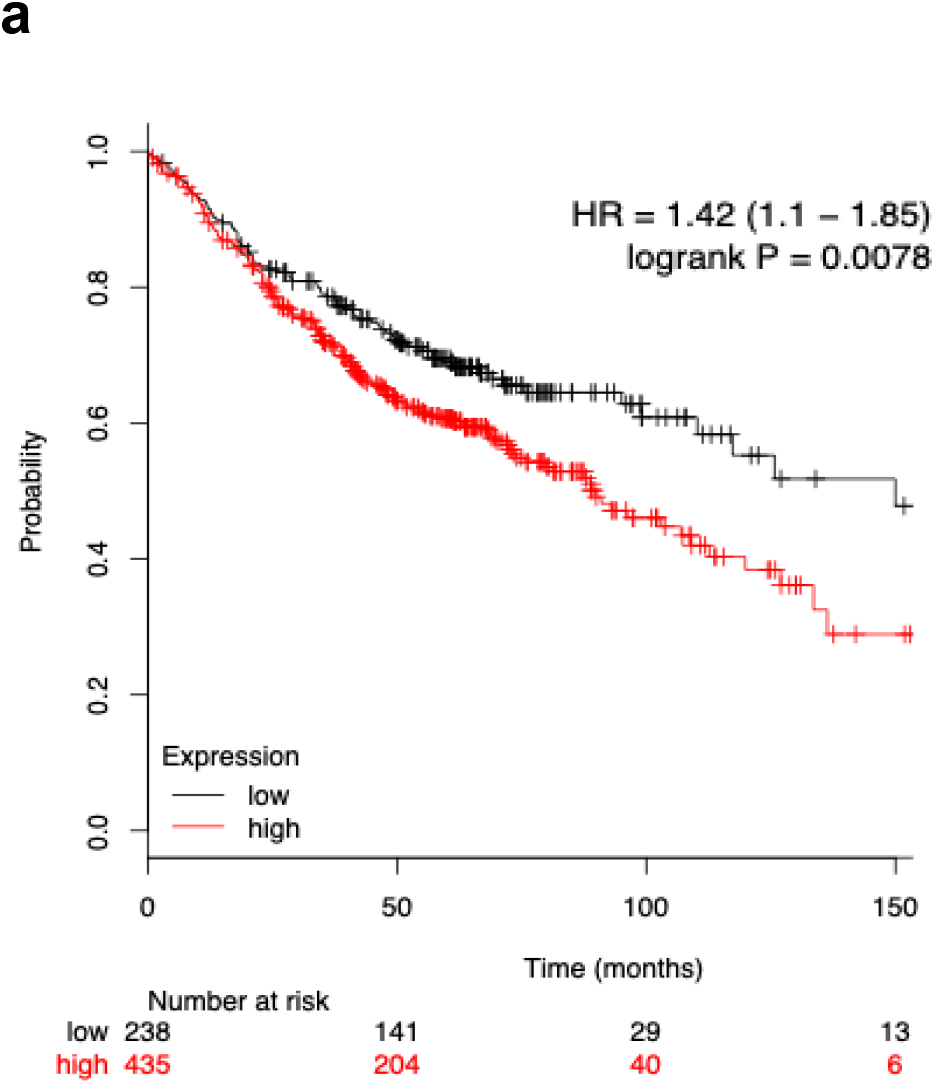
High SREBP1 expression correlates with poor survival in LUAD patients: a) Kaplan-Meier analysis of an integrative lung cancer microarray database showing expression levels of SREBP1 transcript (red:high, black:low) and association with overall survival. Data obtained from Km-plotter website. *p<0.05.

